# Validation of the NovaSeq6000 Platform and automated library preparation for CE-IVD equivalence

**DOI:** 10.1101/2025.08.19.671033

**Authors:** Elena Pasquinelli, Giulia Rollo, Flavia Tinella, Miriana Danelli, Samantha Minetto, Giulia Casamassima, Michel Bader, Laura Calonaci, Olga Lorenza Colavecchio, Rossella Tita, Roberta Mancini, Margherita Baldassarri, Caterina Lo Rizzo, Anna Maria Pinto, Francesca Ariani, Chiara Fallerini, Paola Montagna, Vincenzo Mezzatesta, Mirella Bruttini, Alessandra Renieri

## Abstract

The implementation of next-generation sequencing (NGS) technologies in clinical diagnostics requires rigorous validation of sequencing platforms and analytical workflows. In this study, we validated the performance of the Illumina NovaSeq6000 Research Use Only (RUO) platform, combined with automated library preparation using the Hamilton Microlab STAR system, by comparison to the CE-IVD certified NovaSeq6000Dx platform, which currently relies on manual library preparation. A total of 96 clinical samples underwent whole-exome sequencing (WES) on both platforms. Variant detection performance was assessed for single nucleotide variants (SNVs) and copy number variants (CNVs). The RUO platform demonstrated 100% concordance with the CE-IVD system for clinically relevant SNVs, with full agreement across positive, negative, and overall percent agreement metrics. For CNVs larger than 150 kb, the positive percent agreement was 79%, rising to 91.7% for CNVs >900 kb. Coverage uniformity and autosomal callability were consistently high across platforms. These results confirm the analytical equivalence of the NovaSeq6000 RUO configuration with automated library preparation for clinical-grade WES. This validation framework supports the adoption of scalable, cost-effective workflows that can achieve diagnostic performance comparable to CE-IVD certified systems and may facilitate routine implementation of exome sequencing in clinical laboratories.

**Graphical Abstract:** 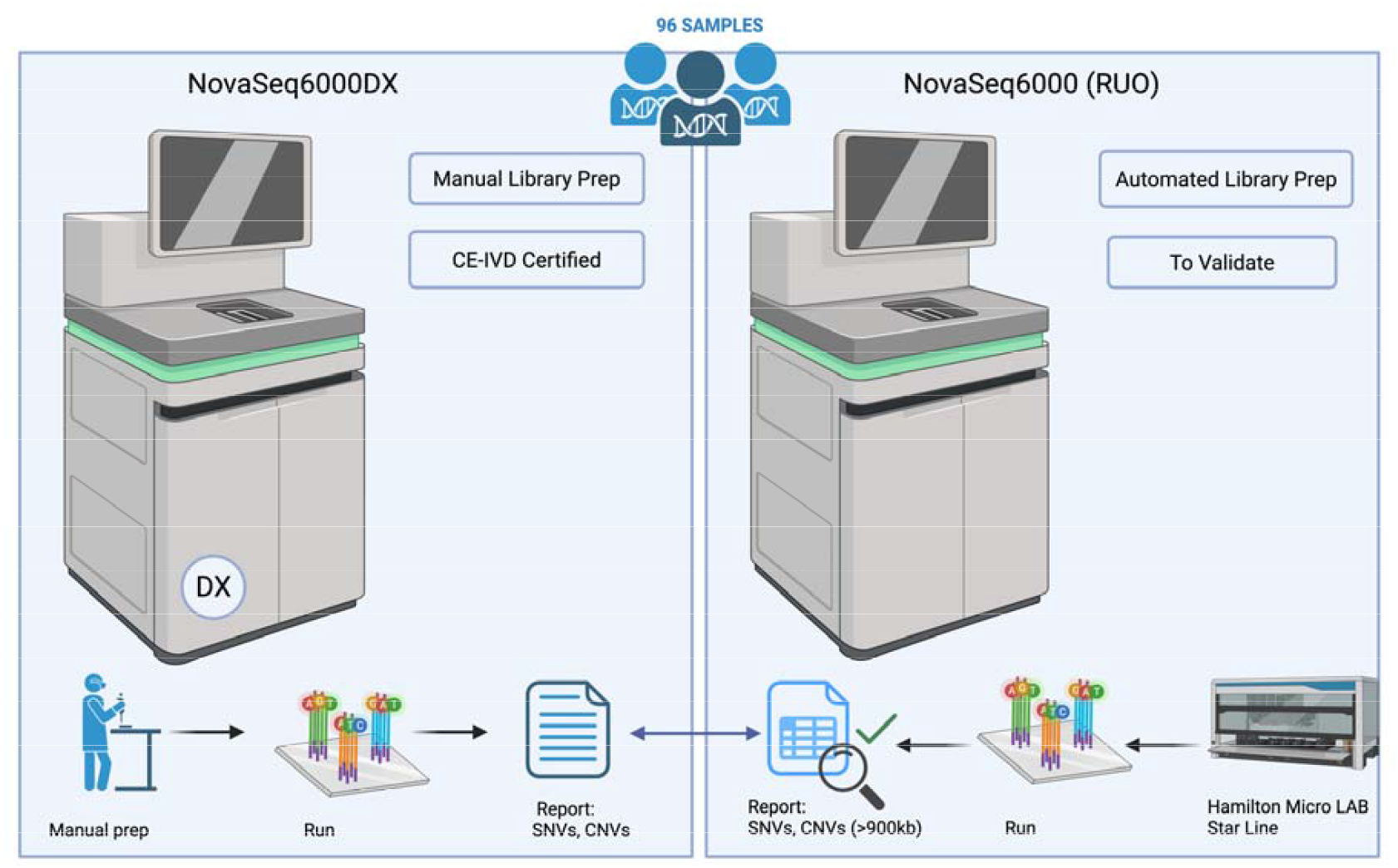

## Introduction

The advent of Next-Generation Sequencing (NGS) technologies has revolutionized the field of clinical genetics, enabling broad-scale genomic investigations that significantly improve the diagnostic yield for patients with suspected genetic disorders. Among the most widely adopted NGS applications in clinical practice is whole exome sequencing (WES), which targets the protein-coding regions of the genome and has proven to be a powerful tool for identifying pathogenic variants in a wide range of Mendelian conditions. However, the implementation of WES for diagnostic purposes requires rigorous validation of both the sequencing platform and the associated laboratory workflows, to ensure compliance with regulatory standards and to guarantee the reliability, accuracy, and reproducibility of results [1].

This procedure arises from the need to internally validate the exome-based diagnostic process established at the Medical Genetics Unit (U.O.C. Genetica Medica) of the Azienda Ospedaliera Universitaria Senese (AOUS) Italy, with particular attention to the use of an internally available sequencing platform that is not CE-IVD marked. Specifically, the NovaSeq6000 Sequencing System by Illumina, does not possess the CE-IVD certification required for in vitro diagnostic use in Europe. Consequently, in light of the recent release of the NovaSeq6000Dx System, an updated CE-IVD certified version of the same platform, it is essential to undertake a comprehensive internal validation process to evaluate the performance characteristics and diagnostic suitability of the existing non-CE-IVD system [2-3].

The validation process also encompasses the automated library preparation system, Hamilton Microlab Star, which plays a critical role in sample processing and contributes to the overall quality and reproducibility of sequencing results [4]. The goal of this technical procedure is to establish standardized operational guidelines for performing a comparative evaluation, risk assessment, and functional verification protocol. This will support the internal validation of the NovaSeq6000 system relative to the NovaSeq Dx model released by Illumina in 2022.

This internal validation initiative is crucial for maintaining high diagnostic standards within the AOUS Medical Genetics Laboratory and for ensuring the continued reliability of genetic test results provided to clinicians and patients. Moreover, the publication of this validation framework may provide valuable guidance to the broader scientific and medical community by offering a reproducible and transparent model for validating WES workflows in clinical settings.

As the use of WES becomes increasingly widespread in diagnostic practice, sharing structured validation approaches contributes to the harmonization of procedures and promotes quality assurance across laboratories.

## Materials and Methods

### 1. Study Design

This validation protocol aims to evaluate the analytical performance of the NovaSeq6000 Sequencing System (Illumina), currently in use at the Medical Genetics Unit of the Azienda Ospedaliera Universitaria Senese (AOUS), by comparing its outputs with those generated by a CE-IVD marked NovaSeq6000Dx system, used in an accredited external Reference Center. The analysis focuses on whole-exome sequencing (WES) across 96 samples, reflecting routine clinical diagnostic scenarios (Figure 1).

**Figure 1.**
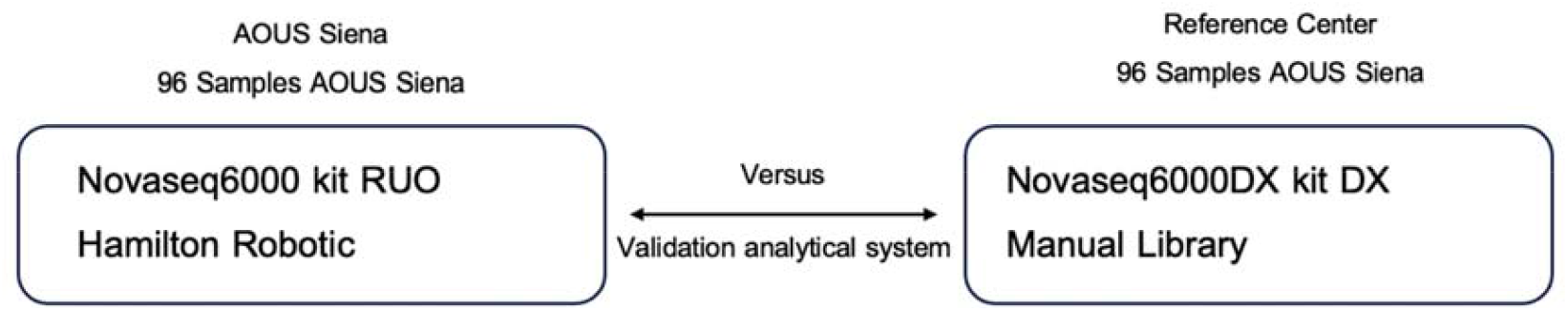
Comparative workflow for analytical validation using 96 AOUS Siena samples. The automated pipeline (left) utilizes the NovaSeq6000 RUO kit with Hamilton robotic library preparation, while the reference pipeline (right) uses the NovaSeq6000DX DX kit with manual library preparation.

### 2. Sample Selection and Composition

A total of 96 samples were selected to reflect realistic diagnostic scenarios encountered in clinical practice. The cohort included both individual and family-based cases. Specifically, 51 unrelated individuals were analyzed, including patients with cancer, alongside 15 family units composed of trios or quads, primarily related to neurodevelopmental disorders. In these cases, the probands were analyzed in conjunction with parental samples, following a TRIO-based study design. All 96 samples originated from the AOUS Medical Genetics Laboratory at AOUS and were processed in parallel on both sequencing platforms using aliquots of the same genomic DNA. This approach ensured that technical variability between runs was minimized, and that observed differences in variant detection were attributable solely to platform-specific performance and not to pre-analytical differences in sample preparation.

### 3. Sequencing Platforms

The platform under validation was the NovaSeq6000 Sequencing System, located at the Medical Genetics Unit of AOUS Siena (Table 1). Although not CE-IVD certified, Illumina has declared that this instrument is technically equivalent in hardware and performance to the NovaSeq6000Dx, the reference platform used for comparison. The NovaSeq6000Dx system is CE-IVD marked and installed in an accredited public or private diagnostic laboratory, serving as the gold standard for validation.

**Table 1.**
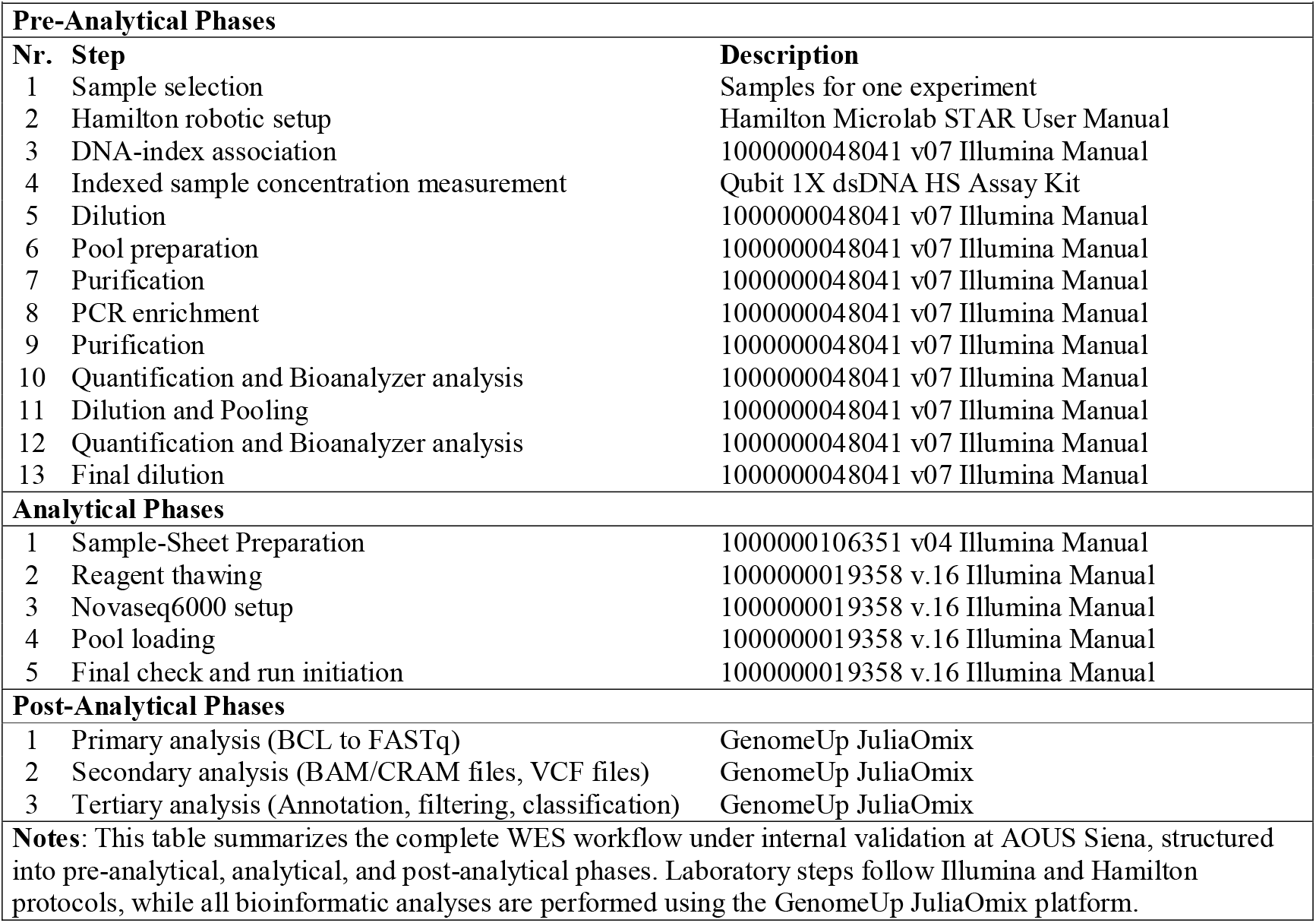
Whole Exome Sequencing Workflow Under Validation at AOUS Siena

At AOUS, sequencing libraries were prepared using the Illumina DNA Prep with Exome 2.5 Enrichment Kit (RUO), Set B for 96 samples (Cat. No. 20077595), combined with the Twist Bioscience Exome 2.5 Panel (Cat. No. 20076914) for target enrichment. Sequencing was performed on the NovaSeq 6000 S2 Reagent Kit v1.5 (300 cycles), RUO (Cat. No. 20028314). Library preparation steps were fully automated using the Hamilton Microlab STAR system (NGS STAR model).

In contrast, the Reference Center employed a manual library preparation workflow using the Illumina DNA Prep with Enrichment Dx with UD Indexes Set A (96 samples, Cat. No. 20051352), paired with the same Twist Exome 2.5 Panel. Sequencing was carried out using the NovaSeq 6000Dx S2 Reagent Kit v1.5 (300 cycles), CE-IVD certified (Cat. No. 20046931).

### 4. Bioinformatic Analysis

Sequencing data generated from both AOUS Siena and the Reference Center underwent standardized bioinformatic using the JuliaOmix platform developed by GenomeUp processing to ensure consistency and minimize technical bias. Primary analysis, including base calling and conversion of BCL to FASTQ files, was performed using the Illumina DRAGEN platform (v4.3.6), which also supported secondary analysis tasks such as alignment to the reference genome (GRCh37), duplicate marking, and variant calling [5]. For comparison and refinement, variant calling was further validated using GATK v4.4 and DeepVariant v1.5, enabling robust detection of single nucleotide variants (SNVs) and small insertions/deletions (indels) [6-7]. These variant sets were harmonized and annotated using Ensembl Variant Effect Predictor (VEP) v113, which provided functional consequences for each variant based on the GRCh37 reference [8].

Comprehensive variant annotation included integration of multiple population and clinical databases. Population allele frequencies were retrieved from both gnomAD v2.1.1 (GRCh37) and v4.1 (GRCh37), while known variant references were queried against dbSNP build 146 and ClinVar (release 20250209) [9-10-11]. Predictive functional scores for missense and splice-altering variants were incorporated from dbNSFP v4.4 and dbscSNV v1.1, supporting variant prioritization and clinical interpretation. Only variants classified as ACMG classes 3, 4, or 5 and relevant to the clinical indication of each patient were retained for comparative analysis. These represent clinically significant findings that would be considered reportable in a real-world diagnostic setting. For each such variant, zygosity, functional annotation, and key quality metrics were documented and summarized in Supplementary Table 1.

Copy number variant (CNV) detection was performed using the Illumina DRAGEN platform (v4.3.6) applied to whole-exome sequencing (WES) data generated with the Illumina Exome 2.5 Panel (GRCh37). CNV segmentation was carried out using the high-sensitivity low-mappability (HSML) mode, with the segmentation window size set to 500 base pairs (-cnv-width-size 500). This setting divided the total target regions (∼262 Mb) into approximately 525,210 windows, enabling genome-wide assessment of copy number profiles at moderate resolution. CNV merging was disabled by default, in accordance with Illumina recommendations for WES analysis, given the spacing between targeted intervals. The merge threshold was maintained at the default linear copy ratio difference of 0.4, allowing for detection of focal CNVs while reducing the risk of artificial fusion of distant segments. For validation purposes, only CNVs classified as clinically relevant and associated with the patient’s diagnostic question were considered. These events would be deemed reportable in a clinical setting, and include deletions or duplications ≥150 kb overlapping known disease-associated loci and are reported in Supplementary Table 2.

### 5. Analytical System Validation Declaration

This validation was qualitative in nature and focused on the presence or absence of expected variants, as determined by a reference table containing 96 pseudonymized samples and their known variant profiles. Sequencing results obtained at AOUS were directly compared to those generated using the CE-IVD certified NovaSeq6000Dx system at an external accredited Reference Center, following Illumina’s standard validation protocol. The assessment of analytical performance included evaluation of accuracy, precision and clinical performance[12]. Accuracy was determined by the concordance of variant calls between the two systems. Precision was measured by the percentage of positive calls (PPC), defined as the number of correctly identified variants over the total number of evaluable variant positions. Clinical performance indicators were also calculated. Positive Percent Agreement (PPA) was defined as the proportion of variant loci correctly reported by the test system. Negative Percent Agreement (NPA) referred to the correct identification of wild-type loci. Overall Percent Agreement (OPA) represented the combined concordance for both variant and wild-type loci. Additionally, the percentage of positive and negative calls (PPC and PNC), excluding low-confidence or low-depth regions, was used to further quantify analytical robustness. The callability of autosomal regions was also evaluated as the proportion of non-N bases in targeted autosomal intervals with valid genotype calls.

## Results

### 1. Run metrics

Run metrics from the two sequencing workflows demonstrated overall comparability in terms of read alignment and depth of coverage. The total number of aligned reads was similar between the two runs, while the average alignment depth and mean coverage across autosomal and sex chromosomes were higher in the AOUS Siena dataset. The percentage of target bases achieving standard depth thresholds (≥15×, ≥20×, ≥50×, and ≥100×) remained consistently high in both datasets, with minimal variation observed across platforms. A slight increase in duplication rate was noted in the AOUS Siena run, consistent with the use of automated library preparation, but without compromising coverage or callability across targeted regions (Table 2 and Figure 2A,B).

**Table 2.** Run Metrics Comparison Between REF Center and AOUS Siena

| Metrics | REF Center | AOUS Siena |
| --- | --- | --- |
| Aligned Reads | 79243944.61 | 76562561.09 |
| Mean Alignment Depth | 124.75 | 139.65 |
| Autosomal Mean Coverage | 125.94 | 140.89 |
| Chr X Mean Coverage | 100.87 | 115.09 |
| Chr Y Mean Coverage | 14.89 | 17.34 |
| Chr M Mean Coverage | 0.0 | 0.0 |
| Number of Marked Duplicate Reads | 8503316.64 | 14338459.55 |
| Percentage of marked duplicate reads | 9.01 | 14.92 |
| Percentage of Bases in target_bed with $\geq 15\times$ Coverage | 98.19 | 97.35 |
| Percentage of Bases in target_bed with $\geq 20\times$ Coverage | 97.88 | 96.58 |
| Percentage of Bases in target_bed with $\geq 50\times$ Coverage | 91.11 | 86.39 |
| Percentage of Bases in target_bed with $\geq 100\times$ Coverage | 60.27 | 60.49 |

**Figure 2.**
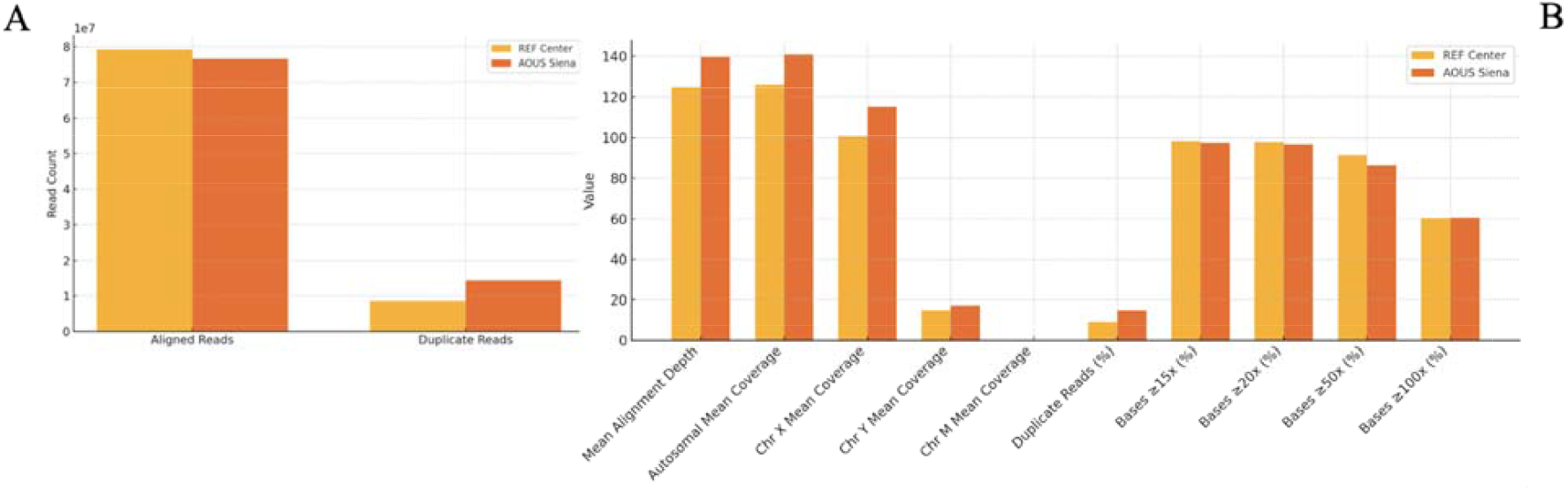
Comparative analysis of sequencing run metrics between AOUS Siena and the Reference Center. (A) Total number of aligned reads and number of marked duplicate reads for each sequencing run. (B) Percentage metrics for coverage thresholds (≥15×, ≥20×, ≥50×, and ≥100×) and percentage of marked duplicate reads. The data illustrate comparable sequencing performance between the two centers, with slight variations in duplication rates and coverage depths.

### 2. SNV Validation

To assess the analytical validity of single nucleotide variant (SNV) detection, we compared the results obtained from the NovaSeq6000 system at AOUS Siena with those from the CE-IVD certified NovaSeq6000Dx platform used in the Reference Center. A complete list of detected SNVs for each sample, including genotype concordance, is provided in Supplementary Table 1. Out of 35 clinically relevant samples, all variants identified at the Reference Center were also correctly detected by the AOUS platform. This resulted in a Positive Percent Agreement (PPA) of 100.00%.

Among 61 samples classified as negative by the Reference Center, all were correctly confirmed as negative by the AOUS system, yielding a Negative Percent Agreement (NPA) of 100.00%.

Combining these results, the Overall Percent Agreement (OPA) including both positive and negative concordant calls was 100.00%, underscoring the high reliability of the test workflow under real diagnostic conditions (Figure 3).

**Figure 3.**
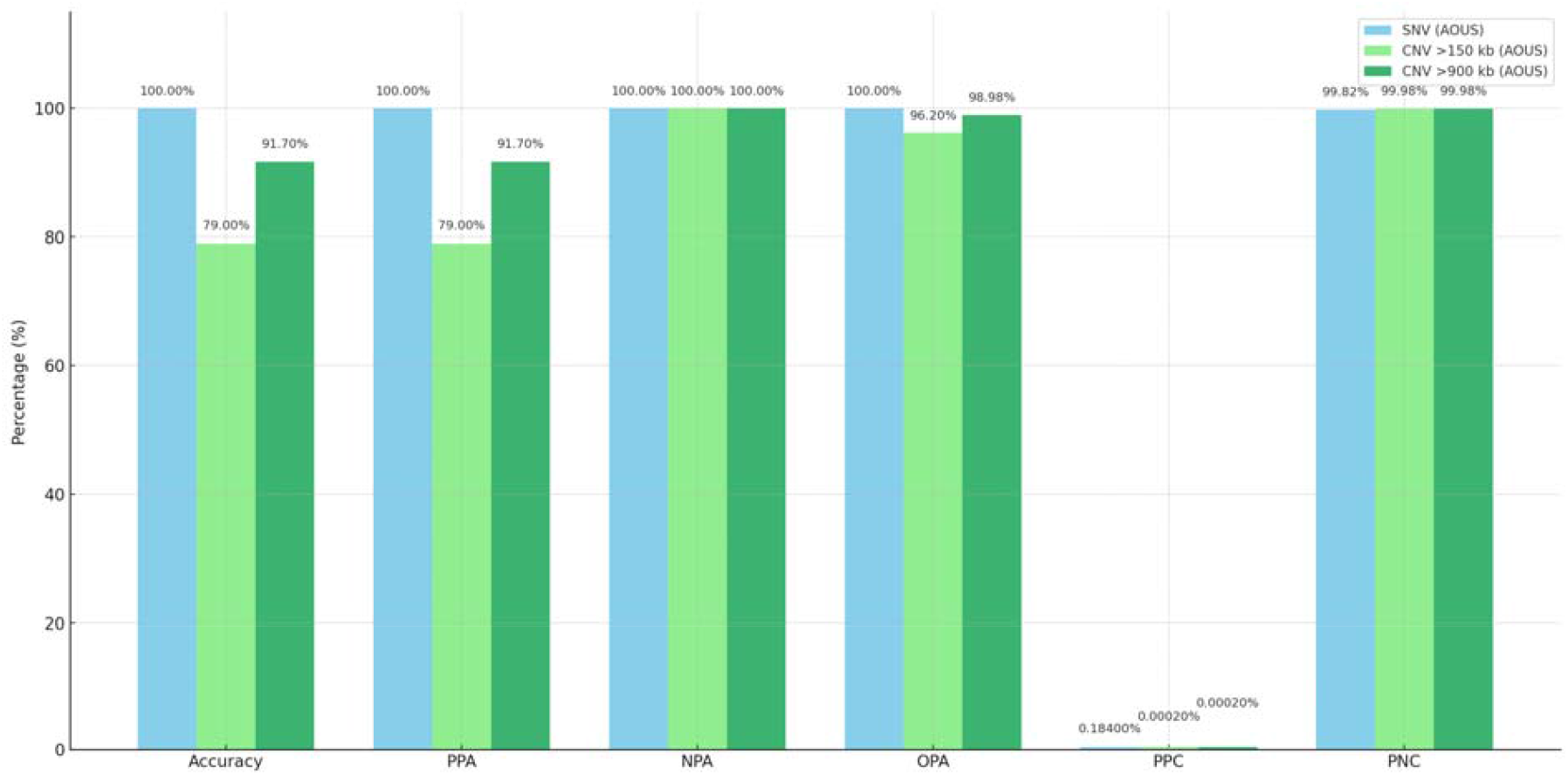
Comparative performance metrics for SNV and CNV detection. Bar chart comparing key analytical and clinical validation parameters: Accuracy, Positive Percent Agreement (PPA), Negative Percent Agreement (NPA), Overall Percent Agreement (OPA), Percent Positive Calls (PPC), and Percent Negative Calls (PNC) across three variant types: single nucleotide variants (SNVs), copy number variants (CNVs) >150 kb, and CNVs >900 kb. Each bar represents the percentage achieved by the AOUS NovaSeq6000-based workflow for the respective metric. While all categories reached 100% NPA, a gradual improvement in PPA and OPA is observed from CNVs >150 kb to CNVs >900 kb, reflecting improved detectability with increasing variant size. SNV performance was uniformly optimal across all metrics.

Analytical performance metrics calculated over all tested base positions (n = 36,547,587) showed a Percent Positive Calls (PPC) of 0.181% and a Percent Negative Calls (PNC) of 99.8187% for the AOUS platform. These values were comparable to those obtained on the reference NovaSeq6000Dx system, indicating a consistent distribution of variant and wild-type calls between the two workflows. The callability of autosomal regions at ≥20× coverage was 96.58%, slightly below the 97.88% obtained on the reference platform.

### 3. CNV Validation (>150kb)

As part of the extended analytical validation, copy number variant (CNV) detection was evaluated for events larger than 150 kb. A total of 96 samples were evaluated for copy number variant (CNV) detection on both the NovaSeq6000Dx (Reference Center) and the NovaSeq6000 RUO platform (AOUS).

The reference center identified 19 clinically relevant CNVs, of which 15 were correctly detected by AOUS, resulting in an accuracy and Positive Percent Agreement (PPA) of 79%.

The AOUS system classified 88 samples as negative, but only 86 of these were true negatives, indicating two false negative results. Full CNV call data, including discordant events, are shown in **Supplementary Table 2**. The resulting Negative Percent Agreement (NPA) was 100%, calculated based on the 86 confirmed negatives over 86 expected negatives. The Overall Percent Agreement (OPA) across the entire cohort was 96.20% (Figure 3). The Percent Positive Calls (PPC) and Percent Negative Calls (PNC) were calculated globally across all CNV calls on the AOUS platform, based on 525,210 genomic segments. PPC was 0.0002% and PNC was 99.98%. The callability of autosomal regions at ≥20× coverage was 96.58% for AOUS, compared to 97.88% on the reference platform.

### 4. CNV Validation (>900kb)

Among the 96 samples analyzed, the reference center identified 12 CNVs larger than 900 kb, of which 11 were correctly detected by the AOUS platform. This corresponds to a Positive Percent Agreement (PPA) and analytical accuracy of 91.7%. The AOUS system classified 93 samples as negative, all of which were also negative according to the reference platform, yielding a Negative Percent Agreement (NPA) of 100% for CNVs larger than 900 kb. One CNV >900 kb was not identified by the AOUS workflow, resulting in a false negative. The Overall Percent Agreement (OPA) between the two systems for this subset was 98.98% (Figure 3). As noted above, PPC and PNC values (0.0002% and 99.98%, respectively) were calculated across all CNV calls, including those in this >900 kb subset, based on the 525,210 genomic segments evaluated. The callability of autosomal regions at ≥20× coverage remained at 96.58% for the AOUS platform, versus 97.88% on the reference system.

## DISCUSSION

This study demonstrates that the NovaSeq6000 platform, combined with an automated library preparation workflow, achieves excellent analytical performance that is largely equivalent to the CE-IVD certified NovaSeq6000Dx system.

This study focused on the analytical validation of SNVs and CNVs detectable through a WES-based pipeline. Other classes of structural variants (SVs), such as inversions, translocations, and complex insertions, were not assessed, as these typically require whole-genome sequencing or long-read technologies for accurate resolution.

For single nucleotide variants (SNVs), the Positive Percent Agreement, Negative Percent Agreement, and Overall Percent Agreement all reached 100%, underscoring the capacity of the workflow to reliably identify clinically relevant variants. This is consistent with previously published studies reporting high concordance rates (>99%) between RUO and CE-IVD configurations of Illumina platforms [13].

For copy number variants (CNVs) larger than 150 kb, the Positive Percent Agreement was 79%, rising to 91.7% for events exceeding 900 kb, and the Overall Percent Agreement approached 99%. The observed Positive Percent Agreement (PPA) of 79% for CNVs larger than 150 kb may appear moderate for clinical validation. However, this value aligns with expectations for CNV detection from whole-exome sequencing (WES) data, where sensitivity typically ranges from 75% to 90%, depending on event size, genomic context, and bioinformatic strategy [14,15].

Prior studies have demonstrated that detection sensitivity increases significantly for CNVs ≥100 kb, as smaller events often fall below the resolution limits of exon-level read-depth analysis due to sparse target coverage and increased signal variability [16]. While the literature supports 100 kb as a general lower boundary, we adopted a more conservative threshold of 150 kb to minimize false negatives and prioritize events with greater diagnostic impact. This choice reflects a balance between analytical reliability and clinical utility.

Furthermore, sensitivity increased to 91.7% for CNVs exceeding 900 kb, which are more robustly detected due to clearer coverage signals and better alignment performance. These findings underscore the appropriateness of our validation framework for capturing clinically reportable CNVs with meaningful diagnostic impact. These large events typically generate more robust and uniform read-depth signals, enhancing detection confidence in WES-based pipelines. The rationale for selecting the 900□ kb threshold is supported by prior evidence from array-based studies, which demonstrated sensitivity and specificity values of 90–95% for genic CNVs ≥900□kb, both deletions and duplication [17].

These results reflect the known limitations of exome-based CNV detection, which tends to perform optimally on large events with clear coverage shifts, while smaller or complex rearrangements remain more challenging to resolve accurately [18,19].

Notably, among the 12 CNVs ≥900 kb identified by the reference center, ten were concentrated in a single sample, consistent with a suspected chromothripsis event. This complex genomic rearrangement was accurately detected by the AOUS pipeline, demonstrating the system’s sensitivity even in the presence of high-structural-load cases.

No additional orthogonal validation was performed on discordant CNV calls because the CE-IVD certified NovaSeq6000Dx platform served as the reference method. Given its regulatory approval and use in an accredited diagnostic center, the CE-IVD workflow was considered an appropriate benchmark for analytical comparison. This approach is consistent with prior studies that have used clinically validated platforms as reference comparators for assessing research-use-only (RUO) systems, particularly when aiming to assess equivalence rather than absolute performance [14,15].

From a diagnostic perspective, the overall yield of pathogenic or likely pathogenic findings in this study was approximately 36.5%, which aligns closely with prior large-scale investigations of clinical WES. For example, Farwell et al. (2015) reported a diagnostic yield of 26% in unselected cohorts, while more recent studies have achieved rates approaching 40% in carefully phenotyped cases with suspected monogenic disorders [20,21]. These comparisons support the conclusion that the validated pipeline not only demonstrates analytical equivalence but also maintains high diagnostic utility.

Evaluating run quality metrics, we observed that the AOUS platform had a higher duplication rate (14.92%) compared to the CE-IVD system (9.01%). This difference did not affect variant detection, as overall coverage depth, callability, and variant concordance remained fully comparable. Slightly elevated duplication rates are commonly observed in automated workflows, particularly when using robotic liquid handlers for library preparation. These values remain within acceptable quality control thresholds for clinical WES and have been reported in previous high-throughput studies using similar setups [19,20]. Therefore, the observed duplication rate does not compromise the analytical robustness or diagnostic reliability of the AOUS pipeline.

One limitation of this study is the absence of intra-run or inter-run replicates, which are important for assessing technical precision and reproducibility over time. While the comparative design effectively demonstrated analytical equivalence with a CE-IVD system, future validation phases may incorporate repeated measures across multiple runs to further strengthen the internal quality control framework.

The internal validation framework we applied here provides evidence that a fully automated WES workflow based on the NovaSeq6000 can be effectively implemented for routine clinical use without compromising accuracy or sensitivity. Compared to the CE-IVD system, this configuration offers substantial advantages in scalability and cost containment, as it allows high-throughput processing without requiring manual library preparation. Furthermore, the ability to deliver diagnostic performance comparable to CE-IVD workflows suggests that laboratories operating under accreditation standards may consider similar validation strategies to optimize resource utilization while maintaining regulatory compliance.

## CONCLUSION

The NovaSeq6000 demonstrated excellent analytical performance in detecting clinically relevant SNVs and high reliability for large CNVs, supporting its suitability for diagnostic WES in clinical setting. This work validates the entire analytical workflow by integrating the NovaSeq6000 sequencing platform with automated library preparation using the Hamilton system, offering a fully automated and scalable solution that is more suitable for high-throughput clinical use than the NovaSeq6000Dx CE-IVD system, which currently relies on manual library preparation.

Moreover, validating the NovaSeq6000 allows clinical laboratories to achieve diagnostic performance comparable to DX-certified platform without compromising diagnostic accuracy and saving costs. This validation framework provides a model for harmonizing WES workflows in routine clinical genetics.

## Data Availability Statement

The raw sequencing data generated in this study cannot be publicly shared due to restrictions in participant informed consent, which did not permit deposition in open-access repositories. Data access may be considered upon reasonable request and pending approval by the relevant ethics committee, subject to GDPR and institutional policies.

## Ethics Statement

The study was conducted in accordance with the ethical principles established by the Declaration of Helsinki. All participants provided written informed consent, and the procedures were approved by the institutional ethics committee (Regional Ethics Committee for Clinical Trials of the Tuscany Region – South-East Area).

## ACKNOWLEDGEMENTS

This work is part of annual department strategic project (2025).

## AUTHOR CONTRIBUTIONS

Conceptualization, E.P., A.R.; methodology, G.R.,G.C., S.M.; C.F., F.A.,O.L.C, LC; software, E.P.,S.M., F.T, M.D.; validation, G.C.,M.B; formal analysis G.R., S.M., E.P., G.C., C.F investigation, E.P., A.R., G.C, G.R., R.D.: resources, A.R.; data curation, E.P., C.F., A.R writing—original draft preparation, E.P.,A.R writing—review and editing E.P., A.R.; ; clinical part of the study and provided patient samples, M.B., A.P.,C.L.,A.R.; visualization, E.P. A.R.; supervision, A.R.; project administration, A.R; funding acquisition, A.R. All authors have read and agreed to the published version of the manuscript.

## SUPPLEMENTARY INFORMATION

Supplementary Table 1

Supplementary Table 2

## References

1. Richards S, Aziz N, Bale S, et al. Standards and guidelines for the interpretation of sequence variants: a joint consensus recommendation of the American College of Medical Genetics and Genomics and the Association for Molecular Pathology. Genet Med. 2015;17(5):405–424. doi:10.1038/gim.2015.30

2. Illumina Inc. NovaSeq 6000Dx Instrument Reference Guide. Document No. 1000000155296 v02; 2022.

3. European Parliament and Council. Regulation (EU) 2017/746 on in vitro diagnostic medical devices. Off J Eur Union. 2017;L117:176–332.

4. Hamilton Company. Microlab STAR User Manual. Hamilton Robotics.

5. Illumina Inc. DRAGEN Bio-IT Platform User Guide v4.3.6. Available from: https://support.illumina.com

6. Poplin R, Chang PC, Alexander D, et al. A universal SNP and small-indel variant caller using deep neural networks. Nat Biotechnol. 2018;36(10):983–987.

7. McKenna A, Hanna M, Banks E, et al. The genome analysis toolkit: a MapReduce framework for analyzing next-generation DNA sequencing data. Genome Res. 2010;20(9):1297–1303.

8. Ensembl Variant Effect Predictor (VEP). Ensembl v113. Available from: https://www.ensembl.org/info/docs/tools/vep/index.html

9. Genome Aggregation Database (gnomAD) v2.1.1 & v4.1. Available from: https://gnomad.broadinstitute.org

10. ClinVar. Clinical Variant Database. NCBI. Release 20250209. Available from: https://www.ncbi.nlm.nih.gov/clinvar

11. Sherry ST, Ward MH, Kholodov M, et al. dbSNP: the NCBI database of genetic variation. Nucleic Acids Res. 2001;29(1):308–311.

12. Clinical and Laboratory Standards Institute (CLSI). MM17-A: Verification and Validation of Multiplex Nucleic Acid Assays. Wayne, PA: CLSI; 2008.

13. Illumina Inc. DRAGEN CNV Pipeline User Guide v4.2; 2022. Available from: https://support.illumina.com

14. Riggs ER, Andersen EF, Cherry AM, et al. Technical standards for the interpretation and reporting of constitutional copy-number variants. Genet Med. 2020;22(2):245–257. doi:10.1038/s41436-019-0686-8

15. Gross AM, Ajay SS, Rajan V, et al. Copy-number variants in clinical genome sequencing: deployment and interpretation for rare and undiagnosed disease. Genet Med. 2019;21(5):1121–1130. doi:10.1038/s41436-018-0282-5

16. Yao R, Zhang C, Yu T, Li N, Hu X, Wang X, et al. Evaluation of three read-depth based CNV detection tools using whole-exome sequencing data. BMC Bioinformatics. 2020;21:97. doi:10.1186/s12859-020-3421-1

17. Zhang Y, Guo Y, Choi H, Glessner J, Hakonarson H, et al. Detecting large copy number variants using exome genotyping arrays in a large Swedish schizophrenia sample. PLoS One. 2014;9(3):e94003. doi:10.1371/journal.pone.0094003

18. Stark Z, Tan TY, Chong B, et al. A prospective evaluation of whole-exome sequencing as a first-tier molecular test in infants with suspected monogenic disorders. Genet Med. 2016;18(11):1090–1096. doi:10.1038/gim.2016.1

19. Clark MM, Stark Z, Farnaes L, et al. Meta-analysis of the diagnostic and clinical utility of genome and exome sequencing and chromosomal microarray in children with suspected genetic diseases. NPJ Genom Med. 2018;3:16. doi:10.1038/s41525-018-0053-8

20. Farwell KD, Shahmirzadi L, El-Khechen D, et al. Enhanced utility of family-centered diagnostic exome sequencing with inheritance model-based analysis: results from 500 unselected families with undiagnosed genetic conditions. Genet Med. 2015;17(7):578–586. doi:10.1038/gim.2014.154

21. Wright CF, FitzPatrick DR, Firth HV. Paediatric genomics: diagnosing rare disease in children. Nat Rev Genet. 2018;19(5):253–268. doi:10.1038/nrg.2017.116

