## Supplementary Table 1-2 for "Validation of the NovaSeq6000 Platform and automated library preparation for CE-IVD equivalence": Supplementary Table_1.pdf

| | Results in reference center | | | | | Results in test center | | | | | Concordance |
|---|---|---|---|---|---|---|---|---|---|---|---|
| **Code** | Reported Gene | c.DNA | Protein | ACMG Class | Zygosity | Reported Gene | c.DNA | Protein | ACMG Class | Zygosity | |
| **Single samples** | | | | | | | | | | | |
| **156047** | SEC23B | c.325G>A | p.Glu109Lys | 5 | heterozygous | SEC23B | c.325G>A | p.Glu109Lys | 5 | heterozygous | 1 |
| **156072** | negative | | | | | negative | | | | | 1 |
| **156074** | MSH2 | c.2519_2530del | p.Val840_Cys843del | 4 | heterozygous | MSH2 | c.2519_2530del | p.Val840_Cys843del | 4 | heterozygous | 1 |
| **156069** | negative | | | | | negative | | | | | 1 |
| **156197** | GLA | c.335G>A | p.Arg112His | 5 | hemizygous | GLA | c.335G>A | p.Arg112His | 5 | hemizygous | 1 |
| **156326** | BRIP1 | c.2087C>T | p.Pro696Leu | 3 | heterozygous | BRIP1 | c.2087C>T | p.Pro696Leu | 3 | heterozygous | 1 |
| **136841** | ERCC2 | c.1308-2A>G | / | 5 | heterozygous | ERCC2 | c.1308-2A>G | / | 5 | heterozygous | 1 |
| **94999** | MYBPC3 | c.1102_1104del | p.Lys368del | 3 | heterozygous | MYBPC3 | c.1102_1104del | p.Lys368del | 3 | heterozygous | 1 |
| **155591** | negative | | | | | negative | | | | | 1 |
| **138168** | FLNC | c.3932C>T | p.Thr1311Ile | 3 | heterozygous | FLNC | c.3932C>T | p.Thr1311Ile | 3 | heterozygous | 1 |
| **79718** | negative | | | | | negative | | | | | 1 |
| **45953** | negative | | | | | negative | | | | | 1 |
| **121721** | ERCC3 | c.325C>T | p.Arg109Ter | 5 | heterozygous | ERCC3 | c.325C>T | p.Arg109Ter | 5 | heterozygous | 1 |
| **100440** | MUTYH | c.1187G>A | p.Gly396Asp | 5 | heterozygous | MUTYH | c.1187G>A | p.Gly396Asp | 5 | heterozygous | 1 |
| **5211-19** | MUTYH | c.1420C>T | Arg474Cys | 3 | heterozygous | MUTYH | c.1420C>T | p.Arg474Cys | 3 | heterozygous | 1 |
| **156428** | MLH3 | c.1119_1122del | p.Phe374MetfsTer8 | 4 | heterozygous | MLH3 | c.1119_1122del | p.Phe374MetfsTer8 | 4 | heterozygous | 1 |
| **156337** | HOXB13 | c.251G>A | p.Gly84Glu | 3 | heterozygous | HOXB13 | c.251G>A | p.Gly84Glu | 3 | heterozygous | 1 |
| **23654** | negative | | | | | negative | | | | | 1 |
| **155895** | negative | | | | | negative | | | | | 1 |
| **155381** | negative | | | | | negative | | | | | 1 |
| **105997** | negative | | | | | negative | | | | | 1 |
| **14279** | negative | | | | | negative | | | | | 1 |
| **156551** | negative | | | | | negative | | | | | 1 |
| **156662** | negative | | | | | negative | | | | | 1 |
| **156674** | negative | | | | | negative | | | | | 1 |
| **152018** | negative | | | | | negative | | | | | 1 |
| **156065** | negative | | | | | negative | | | | | 1 |
| **156669** | MUTYH | c.312C>A | p.Tyr104Ter | 5 | heterozygous | MUTYH | c.312C>A | p.Tyr104Ter | 5 | heterozygous | 1 |
| **10631** | negative | | | | | negative | | | | | 1 |
| **156390** | ongoing | | | | | negative | | | | | 1 |
| **156381** | negative | | | | | negative | | | | | 1 |
| **156692** | negative | | | | | negative | | | | | 1 |
| **156543** | negative | | | | | negative | | | | | 1 |
| **156760** | FANCF | c.224del | p.Gly75ValfsTer6 | 4 | heterozygous | FANCF | c.224del | p.Gly75ValfsTer6 | 4 | heterozygous | 1 |
| **86636** | negative | | | | | negative | | | | | 1 |
| **156766** | negative | | | | | negative | | | | | 1 |
| **152492** | negative | | | | | negative | | | | | 1 |
| **32222** | negative | | | | | negative | | | | | 1 |
| **156846** | negative | | | | | negative | | | | | 1 |
| **156900** | negative | | | | | negative | | | | | 1 |
| **154844** | negative | | | | | negative | | | | | 1 |
| **157206** | negative | | | | | negative | | | | | 1 |
| **145474** | MSH6 | c.3557G>A | p.Gly1186Asp | 4 | heterozygous | MSH6 | c.3557G>A | p.Gly1186Asp | 4 | heterozygous | 1 |
| **157079** | negative | | | | | negative | | | | | 1 |
| **157093** | PTCH1 | c.3114C>G | p.Cys1038Trp | 3 | heterozygous | PTCH1 | c.3114C>G | p.Cys1038Trp | 3 | heterozygous | 1 |
| **157174** | CHEK2 | c.1180G>A | p.Glu394Lys | 3 | heterozygous | CHEK2 | c.1180G>A | p.Glu394Lys | 3 | heterozygous | 1 |
| **157129** | negative | | | | | negative | | | | | 1 |
| **88340** | MAP2K1 | c.803C>G | p.Ala268Gly | 3 | heterozygous | MAP2K1 | c.803C>G | p.Ala268Gly | 3 | heterozygous | 1 |
| **157073** | negative | | | | | negative | | | | | 1 |
| **157349** | ATM | c.8581A>G | p.Ile2861Val | 3 | heterozygous | ATM | c.8581A>G | p.Ile2861Val | 3 | heterozygous | 1 |
| **26621** | ERCC5 | c.2751dup | p.Leu918IlefsTer12 | 4 | heterozygous | ERCC5 | c.2751dup | p.Leu918IlefsTer12 | 4 | heterozygous | 1 |
| **35877** | negative | | | | | negative | | | | | 1 |
| **134354** | LDLR | c.1358+2T>C | / | 4 | heterozygous | LDLR | c.1358+2T>C | / | 4 | heterozygous | 1 |
| **157535** | negative | | | | | negative | | | | | 1 |
| **129743** | negative | | | | | negative | | | | | 1 |
| **153186** | negative | | | | | negative | | | | | 1 |
| **150465** | RAD51 | c.776A>C | p.Glu259Ala | 4 | heterozygous | RAD51 | c.776A>C | p.Glu259Ala | 4 | heterozygous | 1 |
| **Trios** | | | | | | | | | | | |
| **4248-23P** | LRP1<br>LRP1 | c.6477C>A<br>c.10824C>A | p.Cys2159Ter<br>p.Asp3608Glu | 4<br>3 | heterozygous | LRP1<br>LRP1 | c.6477C>A<br>c.10824C>A | p.Cys2159Ter<br>p.Asp3608Glu | 4<br>3 | heterozygous | 1 |
| **156178F** | LRP1 | c.10824C>A | p.Asp3608Glu | 3 | heterozygous | LRP1 | c.10824C>A | p.Asp3608Glu | 3 | heterozygous | 1 |
| **156179M** | LRP1 | c.6477C>A | p.Cys2159Ter | 4 | heterozygous | LRP1 | c.6477C>A | p.Cys2159Ter | 4 | heterozygous | 1 |
| **156448P** | negative | | | | | negative | | | | | 1 |
| **156452M** | negative | | | | | negative | | | | | 1 |
| **156453F** | negative | | | | | negative | | | | | 1 |
| **145038P** | negative | | | | | negative | | | | | 1 |
| **145039F** | negative | | | | | negative | | | | | 1 |
| **145040M** | GLA | c.335G>A | p.Arg112His | 5 | heterozygous | GLA | c.335G>A | p.Arg112His | 5 | heterozygous | 1 |
| **145898P** | NF1 | c.3461A>G | p.Asn1154Ser | 3 | heterozygous | NF1 | c.3461A>G | p.Asn1154Ser | 3 | heterozygous | 1 |
| **145968M** | NF1 | c.3461A>G | p.Asn1154Ser | 3 | heterozygous | NF1 | c.3461A>G | p.Asn1154Ser | 3 | heterozygous | 1 |
| **156578F** | negative | | | | | negative | | | | | 1 |
| **156526P** | TECR<br>TECR | c.66+25T>C<br>c.16-45C>G | / | 3<br>3 | heterozygous | TECR<br>TECR | c.66+25T>C<br>c.16-45C>G | / | 3<br>3 | heterozygous | 1 |

| | | | | | | | | | | | |
|---|---|---|---|---|---|---|---|---|---|---|---|
| **156683M** | TECR | c.66+25T>C | / | 3 | heterozygous | TECR | c.66+25T>C | / | 3 | heterozygous | 1 |
| **156747F** | TECR | c.16-45C>G | / | 3 | heterozygous | TECR | c.16-45C>G | / | 3 | heterozygous | 1 |
| **148985P** | negative | | | | | negative | | | | | 1 |
| **148921M** | negative | | | | | negative | | | | | 1 |
| **156951F** | negative | | | | | negative | | | | | 1 |
| **138051P** | negative | | | | | negative | | | | | 1 |
| **80777F** | negative | | | | | negative | | | | | 1 |
| **138329M** | negative | | | | | negative | | | | | 1 |
| **156842P** | MEFV | c.2080A>G | p.Met694Val | 5 | heterozygous | MEFV | c.2080A>G | p.Met694Val | 5 | heterozygous | 1 |
| **156843M** | negative | | | | | negative | | | | | 1 |
| **156852F** | MEFV | c.2080A>G | p.Met694Val | 5 | heterozygous | MEFV | c.2080A>G | p.Met694Val | 5 | heterozygous | 1 |
| **157068P** | LDLR | c.1567G>A | p.Val523Met | 5 | heterozygous | LDLR | c.1567G>A | p.Val523Met | 5 | heterozygous | 1 |
| **157290F** | negative | | | | | negative | | | | | 1 |
| **157229M** | LDLR | c.1567G>A | p.Val523Met | 5 | heterozygous | LDLR | c.1567G>A | p.Val523Met | 5 | heterozygous | 1 |
| **157025P** | negative | | | | | negative | | | | | 1 |
| **157125M** | negative | | | | | negative | | | | | 1 |
| **157127F** | negative | | | | | negative | | | | | 1 |
| **Quartet** | | | | | | | | | | | |
| **153591P** | negative | | | | | negative | | | | | 1 |
| **153593S** | negative | | | | | negative | | | | | 1 |
| **153236M** | negative | | | | | negative | | | | | 1 |
| **156264F** | negative | | | | | negative | | | | | 1 |
| **Quintet** | | | | | | | | | | | |
| **132189P** | negative | | | | | negative | | | | | 1 |
| **120997B1** | negative | | | | | negative | | | | | 1 |
| **21799B2** | negative | | | | | negative | | | | | 1 |
| **156934M** | negative | | | | | negative | | | | | 1 |
| **156931F** | negative | | | | | negative | | | | | 1 |
