## Supplementary Table 1-2 for "Validation of the NovaSeq6000 Platform and automated library preparation for CE-IVD equivalence": Supplementary Table_2 .pdf

| | Results in reference center (Rome) | | | | | | Results in test center (Siena) | | | | | | Concordance |
|---|---|---|---|---|---|---|---|---|---|---|---|---|---|
| **Code** | Location | Size | Disease gene(s) involved | ACMG Class | DEL/DUP | Genotype | Location | Size | Disease gene(s) involved | ACMG Class | DEL/DUP | Genotype | |
| **Single samples** | | | | | | | | | | | | | |
| **156047** | Chr22:22988781:23657664 | 668883 (668 Kb) | BCR | 5 | DEL | Heterozygous | Chr22:22988781:23657664 | 668883 (668 Kb) | BCR | 5 | DEL | Heterozygous | 1 |
| **156072** | chr8:42862989:43218462 | 355473 (355 Kb) | POMK, HGSNAT | 5 | DUP | Heterozygous | chr8:42862989:43218462 | 355473 (355 Kb) | POMK, HGSNAT | 5 | DUP | Heterozygous | 1 |
| **156074** | negative | | | | | | negative | | | | | | 1 |
| **156069** | negative | | | | | | negative | | | | | | 1 |
| **156197** | negative | | | | | | negative | | | | | | 1 |
| **156326** | DUP in chr 1, 2, 3, 4, 5, 6, 7, 8, 13, X | * | * | * | * | * | DUP in chr 2, 3, 4, 5, 6, 8, 13, X | * | * | * | * | * | 1 |
| **136841** | chr8:42862989:43057679 | 194690 (194 Kb) | POMK, HGSNAT | 5 | DUP | Heterozygous | chr8:42862989:43218462 | 355473 (355 Kb) | POMK, HGSNAT | 5 | DUP | Heterozygous | 1 |
| **94999** | negative | | | | | | negative | | | | | | 1 |
| **155591** | chr8:42862989:43057679 | 194690 (194 Kb) | POMK, HGSNAT | 5 | DUP | Heterozygous | chr8:42862989:43218462 | 355473 (355 Kb) | POMK, HGSNAT | 5 | DUP | Heterozygous | 1 |
| **138168** | negative | | | | | | negative | | | | | | 1 |
| **79718** | negative | | | | | | negative | | | | | | 1 |
| **45953** | negative | | | | | | negative | | | | | | 1 |
| **121721** | negative | | | | | | negative | | | | | | 1 |
| **100440** | negative | | | | | | negative | | | | | | 1 |
| **5211-19** | negative | | | | | | negative | | | | | | 1 |
| **156428** | negative | | | | | | negative | | | | | | 1 |
| **156337** | Chr16:17564290:18426519 | 862229 (862 Kb) | XYLT1 | 5 | DEL | Heterozygous | negative | | | | | | 0 |
| **23654** | negative | | | | | | negative | | | | | | 1 |
| **155895** | negative | | | | | | negative | | | | | | 1 |
| **155381** | negative | | | | | | negative | | | | | | 1 |
| **105997** | negative | | | | | | negative | | | | | | 1 |
| **14279** | negative | | | | | | negative | | | | | | 1 |
| **156551** | negative | | | | | | negative | | | | | | 1 |
| **156662** | negative | | | | | | negative | | | | | | 1 |
| **156674** | chr17:18933743:19125677 | 191934 (191 Kb) | GRAP | 4 | DEL | Heterozygous | negative | | | | | | 0 |
| **152018** | negative | | | | | | negative | | | | | | 1 |
| **156065** | negative | | | | | | negative | | | | | | 1 |
| **156669** | negative | | | | | | negative | | | | | | 1 |
| **10631** | negative | | | | | | negative | | | | | | 1 |
| **156390** | chr5:69372845:70242003 | 869158 (869 Kb) | SMN1, SMN2 | 5 | DEL | Heterozygous | chr5:69372845:70242003 | 869158 (869 Kb) | SMN1, SMN2 | 5 | DEL | Heterozygous | 1 |
| **156381** | negative | | | | | | negative | | | | | | 1 |
| **156692** | chr2:41607:1442630 | 1401023 (1.4 Mb) | TPO | 4 | DEL | Heterozygous | chr2:41607:1442630 | 1401023 (1.4 Mb) | TPO | 4 | DEL | Heterozygous | 1 |
| **156543** | negative | | | | | | negative | | | | | | 1 |
| **156760** | negative | | | | | | negative | | | | | | 1 |
| **86636** | negative | | | | | | negative | | | | | | 1 |
| **156766** | negative | | | | | | negative | | | | | | 1 |
| **152492** | negative | | | | | | negative | | | | | | 1 |
| **32222** | negative | | | | | | negative | | | | | | 1 |
| **156846** | negative | | | | | | negative | | | | | | 1 |
| **156900** | negative | | | | | | negative | | | | | | 1 |
| **154844** | negative | | | | | | negative | | | | | | 1 |
| **157206** | negative | | | | | | negative | | | | | | 1 |
| **145474** | negative | | | | | | negative | | | | | | 1 |
| **157079** | negative | | | | | | negative | | | | | | 1 |
| **157093** | negative | | | | | | negative | | | | | | 1 |
| **157174** | negative | | | | | | negative | | | | | | 1 |
| **157129** | negative | | | | | | negative | | | | | | 1 |
| **88340** | negative | | | | | | negative | | | | | | 1 |
| **157073** | negative | | | | | | negative | | | | | | 1 |
| **157349** | negative | | | | | | negative | | | | | | 1 |
| **26621** | negative | | | | | | negative | | | | | | 1 |
| **35877** | negative | | | | | | negative | | | | | | 1 |
| **134354** | ChrY:21205048:27770674 | 6565626 (6.5 Mb) | none | 5 | DEL | hemizygous | ChrY:21205048:27770674 | 6565626 (6.5 Mb) | none | 5 | DEL | hemizygous | 1 |
| **157535** | negative | | | | | | negative | | | | | | 1 |
| **129743** | negative | | | | | | negative | | | | | | 1 |
| **153186** | negative | | | | | | negative | | | | | | 1 |
| **150465** | negative | | | | | | | | | | | | 1 |
| **Trios** | | | | | | | | | | | | | |
| **4248-23P** | negative | | | | | | negative | | | | | | |
| **156178F** | negative | | | | | | negative | | | | | | 1 |
| **156179M** | negative | | | | | | negative | | | | | | 1 |
| **156448P** | negative | | | | | | negative | | | | | | 1 |
| **156452M** | negative | | | | | | negative | | | | | | 1 |
| **156453F** | negative | | | | | | negative | | | | | | 1 |
| **145038P** | negative | | | | | | negative | | | | | | 1 |
| **145039F** | negative | | | | | | negative | | | | | | 1 |
| **145040M** | negative | | | | | | negative | | | | | | 1 |
| **145898P** | negative | | | | | | negative | | | | | | 1 |
| **145968M** | negative | | | | | | negative | | | | | | 1 |
| **156578F** | negative | | | | | | negative | | | | | | 1 |
| **156526P** | negative | | | | | | negative | | | | | | 1 |
| **156683M** | negative | | | | | | negative | | | | | | 1 |
| **156747F** | negative | | | | | | negative | | | | | | 1 |
| **148985P** | negative | | | | | | negative | | | | | | 1 |
| **148921M** | negative | | | | | | negative | | | | | | 1 |
| **156951F** | negative | | | | | | negative | | | | | | 1 |
| **138051P** | negative | | | | | | negative | | | | | | 1 |
| **80777F** | negative | | | | | | negative | | | | | | 1 |
| **138329M** | negative | | | | | | negative | | | | | | 1 |
| **156842P** | negative | | | | | | negative | | | | | | 1 |
| **156843M** | negative | | | | | | negative | | | | | | 1 |
| **156852F** | negative | | | | | | negative | | | | | | 1 |
| **157068P** | negative | | | | | | negative | | | | | | 1 |
| **157290F** | negative | | | | | | negative | | | | | | 1 |
| **157229M** | negative | | | | | | negative | | | | | | 1 |
| **157025P** | negative | | | | | | negative | | | | | | 1 |
| **157125M** | negative | | | | | | negative | | | | | | 1 |
| **157127F** | negative | | | | | | negative | | | | | | 1 |
| **Quartet** | | | | | | | | | | | | | |
| **153591P** | negative | | | | | | negative | | | | | | 1 |
| **153593S** | negative | | | | | | negative | | | | | | 1 |
| **153236M** | negative | | | | | | negative | | | | | | 1 |
| **156264F** | negative | | | | | | negative | | | | | | 1 |
| **Quintet** | | | | | | | | | | | | | |
| **132189P** | negative | | | | | | negative | | | | | | 1 |
| **120997B1** | negative | | | | | | negative | | | | | | 1 |
| **21799B2** | negative | | | | | | negative | | | | | | 1 |
| **156934M** | negative | | | | | | negative | | | | | | 1 |
| **156931F** | negative | | | | | | negative | | | | | | 1 |
